# ODGI: understanding pangenome graphs

**DOI:** 10.1101/2021.11.10.467921

**Authors:** Andrea Guarracino, Simon Heumos, Sven Nahnsen, Pjotr Prins, Erik Garrison

**Author notes:** Contributed equally (Andrea Guarracino, Simon Heumos).

## Abstract

**Motivation:** Pangenome graphs provide a complete representation of the mutual alignment of collections of genomes. These models offer the opportunity to study the entire genomic diversity of a population, including structurally complex regions. Nevertheless, analyzing hundreds of gigabase-scale genomes using pangenome graphs is difficult as it is not well-supported by existing tools. Hence, fast and versatile software is required to ask advanced questions to such data in an efficient way.

**Results:** We wrote ODGI, a novel suite of tools that implements scalable algorithms and has an efficient in-memory representation of DNA pangenome graphs in the form of variation graphs. ODGI supports pre-built graphs in the Graphical Fragment Assembly format. ODGI includes tools for detecting complex regions, extracting pangenomic loci, removing artifacts, exploratory analysis, manipulation, validation, and visualization. Its fast parallel execution facilitates routine pangenomic tasks, as well as pipelines that can quickly answer complex biological questions of gigabase-scale pangenome graphs.

**Availability:** ODGI is published as free software under the MIT open source license. Source code can be downloaded from https://github.com/pangenome/odgi and documentation is available at https://odgi.readthedocs.io. ODGI can be installed via Bioconda https://bioconda.github.io/recipes/odgi/README.html or GNU Guix https://github.com/pangenome/odgi/blob/master/guix.scm.

## 1 Introduction

A pangenome models the full set of genomic elements in a given species or clade (Tettelin *et al*., 2008; Computational Pan-Genomics Consortium, 2018; Eizenga *et al*., 2020b). In contrast to reference-based approaches which relate samples to a single genome, these data structures encode the mutual relationships between all the genomes represented (Ballouz *et al*., 2019). A class of methods to represent pangenomes involves sequence graphs (Hein, 1989; Paten *et al*., 2017) where homologous regions between genomes are compressed into single representations of all alleles present in the pangenome. In sequence graphs, node labels are genomic sequences with edges connecting those nodes. A bidirected sequence graph can represent both strands of DNA. On this model, variation graphs add the concept of paths representing linear DNA sequences as traversals through the nodes of the graph (Garrison *et al*., 2018). For example, a path can be a genome, haplotype, contig, or read.

Pangenome graphs can be constructed by multiple sequence alignment (Lee *et al*., 2002; Grasso and Lee, 2004) or by transitively reducing an alignment between sequences to an equivalent, labeled sequence graph (Kehr *et al*., 2014; Garrison, 2019). Current methods to build these graphs are still under active development (Li *et al*., 2020; Armstrong *et al*., 2020; Garrison *et al*., 2021), but they have largely settled on a common data model, represented in the Graphical Fragment Assembly (GFA) format (GFA Working Group, 2016). This standardization supports the development of a reference set of tools that operate on the pangenome graph model.

Pangenome graphs let us encode any kind of variation, allowing the generation of comprehensive data systems that builds the basis for the analyses of genome evolution. The Human Pangenome Reference Consortium (HPRC) and Telomere-to-Telomere (T2T) consortium (Miga *et al*., 2020; Logsdon *et al*., 2021; Nurk *et al*., 2021; Jarvis *et al*., 2022) have recently demonstrated that high-quality haploid and diploid *de novo* assemblies can be routinely generated from third-generation long read sequencing data. We anticipate that *de novo* assemblies of similar quality will become common, leading to demand for methods to analyze pangenomes.

Although pangenome graphs are data structures of utility to researchers (Computational Pan-Genomics Consortium, 2018; Garrison *et al*., 2018; Baaijens *et al*., 2019; Hickey *et al*., 2020; Sibbesen *et al*., 2021), the scientific community still lacks a toolset capable of operating on gigabase-scale pangenome graphs constructed from whole-genome assemblies. Such an effort began with the VG toolkit (Garrison *et al*., 2018), but its tools do not efficiently handle pangenome graphs presenting complex motifs that result from repetitive sequences. Here we refocus the effort with the Optimized Dynamic Genome/Graph Implementation (ODGI) toolkit, a compatible, but independent pangenome graph interrogation and transformation system specifically implemented to handle the data scales encountered when working with pre-built constructed pangenomes comprising hundreds of haplotype-resolved genomes. ODGI offers a set of standard operations on the variation graph data model (Fig. 1), generalizing “genome arithmetic” concepts, like those found in BEDTools (Quinlan and Hall, 2010), to work on pangenome graphs. Furthermore, it provides a variety of tools for graph visualization, sorting, and liftover projections, all critical to understand and exploit pangenome graphs.

**Fig. 1:**
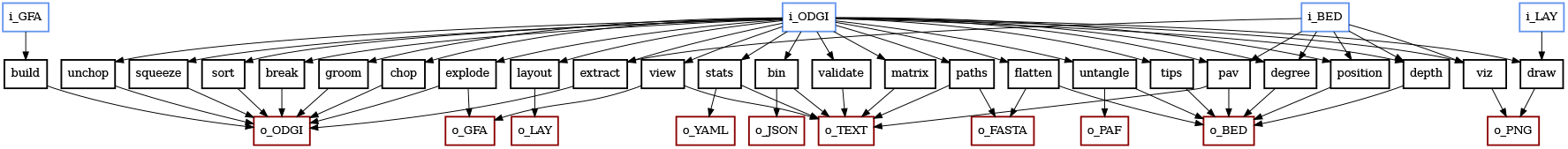
Overview of the methods provided by ODGI (in black) and their supported input (in blue) and output (in red) data formats.

## 2 Model

A pangenome graph is a sequence model that encodes the mutual alignment of many genomes (Garrison, 2019; Eizenga *et al*., 2020b). In the variation graph, *V* = (*N, E, P*), nodes *N* = *n*_1_ … *n*_|*N*|_ contain genomic sequences. Each node *n*_*i*_ has an identifier *i* and an implicit reverse complement 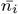, and a node strand *s* corresponds to one of such orientations. Edges *E* = *e*_1_ … *e*_|*E*|_ represent ordered pairs of node strands: *e*_*i*_ = (*s*_*a*_, *s*_*b*_). Paths *P* = *p*_1_ … *p*_|*P*|_ describe walks over node strands: 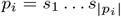. When used as a pangenome graph, *V* expresses sequences, haplotypes, contigs, and annotations as paths. By containing both the sequences and information about their relative variations, the variation graph provides a complete and powerful foundation for many bioinformatic applications.

## 3 Implementation

The ODGI toolkit builds on existing approaches to efficiently store and manipulate pangenome graphs in the form of variation graphs (Garrison *et al*., 2018). Similar to other efficient libraries presenting the HandleGraph model (Eizenga *et al*., 2020a), the implementation of ODGI’s tools rests on three key properties which hold for most pangenome graphs:

1. They are relatively sparse, with low average node degree.
2. They can be sorted so that most edges go between nodes that are close together in the sort order.
3. Their embedded paths are locally similar to each other.

These properties are used to build efficient dynamic variation graph data structures (Siren *et al*., 2020; Eizenga *et al*., 2020a). Sparsity (1) allows us to encode edges *E* using adjacency lists rather than matrices or hash tables. The local linear structure of the graph (2) lets us assign node identifiers that increase along the linear components of the graph, which supports a compact storage of edges and path steps as relativistic (usually small) differences rather than absolute (always large) integer identifiers. Path similarity (3) allows us to write local compressors that reduce the storage cost of collections of path steps.

ODGI improves on prior efforts, based on issues that arose during our work with high-quality *de novo* assemblies that cover almost all parts of the human genome (Logsdon *et al*., 2021; Nurk *et al*., 2021). In particular, we find that it is necessary to support graphs with regions of very high numbers of path traversals (high depth of path coverage of some nodes, the so-called node depth). Such motifs can occur in collapsed structures generated by ambiguous sequence homology relationships in repeats found in the centromeres and other segmental duplications. If we cannot process such regions, we cannot understand them, and our only option is to build graphs that do not include them. Our goal is to build tools that allow for a wide range of uses of pangenome graphs, including cases with potentially high path depth. To seamlessly represent such difficult regions, we followed an approach implemented in the dynamic version of the Graph BWT (GBWT) (Siren *et al*., 2020) and built a node-centric, dynamic, compressed model of the paths. This design supports node-local modification and update of the graph, which lets us build and modify the graph and its paths in parallel.

We store the graph in a vector of node structures, each of which presents a node-local view of the graph sequence, topology, and path layout (Algorithm 1). Expressed in terms of the variation graph *V*, ODGI’s core *Node* structure includes a decoder that maps the neighbors of each node to a dense range of integers. For a given *Node*_*i*_ and neighbor *Node*_*j*_, the decoder itself does not store the *id* of *Node*_*j*_, but rather a compact representation of the relative difference between the node ids: *δ* = *Node*_*i*_.*id* − *Node*_*j*_ .*id*. This keeps the size of the encoding small, per common pangenome graph property (2). We define the edges and path steps traversing the node in terms of this alphabet of *δ*’s.

### Algorithm 1 ODGI’s relativistically-packed *Node* structure and the *Step* structure used to represent the paths as doubly-linked lists.

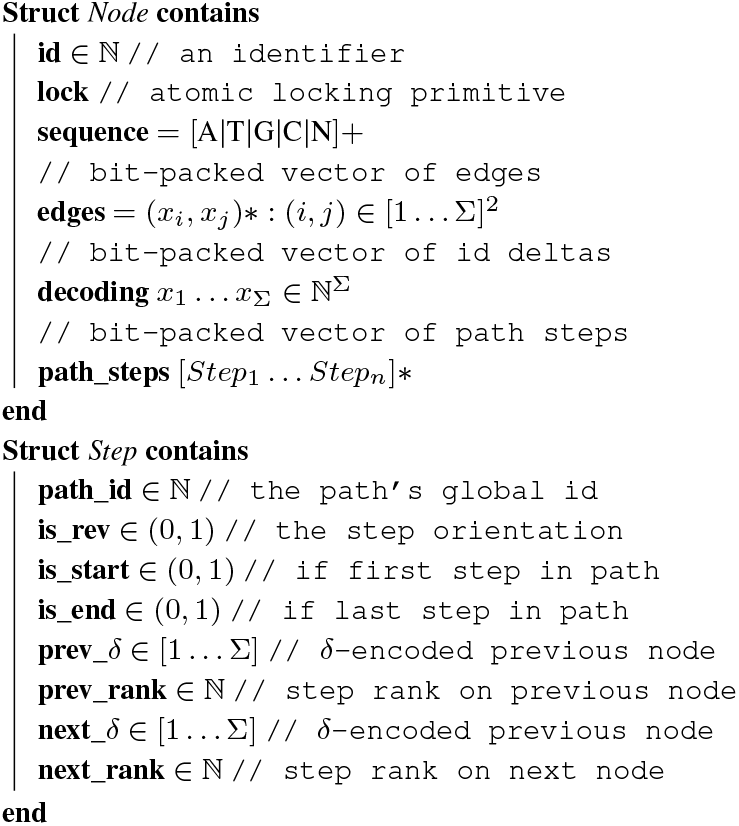

Each structure contains the sequence of the node (*Node*_*i*_.*sequence*), its edges in both directions (*Node*_*i*_.*edges*), and a vector of path steps that describes the previous and next steps in paths that walk across the node (*Node*_*i*_.*path*_*steps*). For efficiency, *Node*_*i*_.*sequence* is stored as a plain string, while the *edges* and *path*_*steps* are stored using a dynamic succinct integer vector that requires *O*(2*nw*) bits for the edges and *O*(5*nw*) bits for the path steps, where *n* is the number of steps on the node and *w* is *≈ log*_2_(*n*) (Prezza, 2017).

To allow edit operations in parallel, each node structure includes a byte-width mutex *lock*. All changes on the graph can involve at most two *Node* structs at a time (both edge and path step representations are doubly-linked). To avoid deadlocks, we acquire the node locks in ascending *Node*.*id* order and release them in descending order. In addition to node-local features of the graph, we must maintain some global information. Specifically, we record the start and end of paths, as well as a name to path id mapping in lock-free hash tables. The use of lock-free hash tables lets us avoid a global lock when looking up path or graph metadata, which would quickly become a bottleneck during parallel operations on the graph. By avoiding global locks, we implement many of the operations in ODGI using maximum parallelism available. This approach is key to enable our methods to scale to the largest pangenome graphs that we can currently build (with hundreds of vertebrate genomes).

## 4 Overview

ODGI provides a set of interrogative and manipulative operations on pangenome graphs. We have established these tools to support our exploration of graphs built from hundreds of large eukaryotic genomes. ODGI’s tools are practical and able to work with high levels of graph complexity, even with regions where paths present very high depth nodes (10^5^ to 10^6^-fold depth). ODGI covers common operations that we have found to be essential when working with complex pangenome graphs:

- *odgi build* constructs the ODGI data model from GFA file (§4.1).
- *odgi view* converts the ODGI data model into GFA file (§4.1).
- *odgi viz* provides a linear visualization of the graph (§5.1).
- *odgi draw* renders a 2D image of the graph (§5.1).
- *odgi extract* excerpts subsets of the graph based on path ranges (§S.3).
- *odgi explode* breaks the graph into connected components (§S.3).
- *odgi squeeze* unifies disjoint graphs (§S.3).
- *odgi chop* breaks long nodes into shorter ones (§S.3).
- *odgi unchop* combines unitig nodes (§S.3).
- *odgi break* removes cycles in the graph (§S.3).
- *odgi prune* removes complex regions (§S.3).
- *odgi groom* resolves spurious inverting links (§S.3).
- *odgi position* lifts coordinates between path and graph positions (§5.2).
- *odgi untangle* deconvolutes paths relative to a reference (§5.2).
- *odgi tips* finds path end points relative to a reference (§S.2).
- *odgi sort* orders the graph nodes (§5.3).
- *odgi layout* establishes a 2D layout (§5.3).
- *odgi matrix* derives the pangenome matrix (§S.5).
- *odgi paths* lists and extracts paths in FASTA (§S.5).
- *odgi flatten* converts the graph to FASTA and BED (§S.5).
- *odgi pav* computes presence-abscence variations (§S.5).
- *odgi stats* provides numerical properties of the graph (§5.4).
- *odgi bin* generates a summarized view of the graph (§S.5).
- *odgi depth* describes node depth over graph and path positions (§5.4).
- *odgi degree* describes node degree over graph and path positions (§5.4).

Each tool focuses on a small set of related operations. Most read or write the native ODGI format (‘og’ extension) (Figure 1) and work with standard text based data formats common to bioinformatics. This supports the implementation of flexible and composable graph processing pipelines based on graphs (GFA/ODGI) and standard bioinformatic data types representing positions, genomic ranges (BED), and pairwise mappings (PAF). We use variation graph paths to provide a universal coordinate system, representing annotations and pairwise sequence relationships using the paths as reference and query sequences. Thus, ODGI provides a set of interfaces that let us approach these graphs from the perspective of standard reference- and sequence-based data models. Indeed, by considering all paths in the graph as potential reference or query sequence, we make graphs invisible to downstream tools that operate on collections of sequences or rely on a reference sequence (*e*.*g*. SAMtools (Li *et al*., 2009)), enabling interoperability. This approach benefits from the information in the graph without requiring that we build an entirely new set of bioinformatic methods to work in this difficult new pangenomic research context.

### 4.1 Building the ODGI model

ODGI maintains its own efficient binary format for storing graphs on disk. We begin by transforming the storage model of the standard GFAv1 (GFA Working Group, 2016) format (in which nodes, edges, and paths are described independently) into the ODGI node-centric encoding with *odgi build*. This construction step can be a significant bottleneck, in particular as the size of the path set of the graph increases. The process itself is lossless. A graph in ODGI format represents everything that is in the input GFAv1 graph, without any loss of information. ODGI does not natively support GFAv2 or rGFA. GFAv2 is similar to GFAv1, but includes process-related annotations of assembly graphs not relevant for pangenome analyses. rGFA embeds a single coordinate hierarchy over the graph that links all sequences into a single base reference genome. This positional model depends on a particular graph induction algorithm *Li et al*. (2020). In contrast, ODGI implements coordinate translation dynamically (e.g. *odgi position* and *odgi untangle*), allowing use of any embedded genome as a reference. Its input graphs can represent any kind of alignment between the genomes. GFAv1 is fully capable of representing many reference genome coordinate systems simultaneously, which supports a reference-agnostic approach that uses the entire pangenome sequence space as a reference system. In doing so, our approach has the advantage of maintaining backward compatibility with existing tools based on genome sequences.

The ODGI data structure (Algorithm 1) allows algorithms that build and modify the graph to operate in parallel, without any global locks. In *odgi build*, we initially construct the node vector in a serial operation that scans across the input GFA file. Then, we serially add edges in the *Node*.*edges* vectors of pairs of nodes. Finally, we create paths in serial, and extend them in parallel by obtaining the mutex *Node*.*lock* for pairs of nodes and by adding the path step in their *Node*.*path*_*steps* vectors. This parallelism speeds ODGI model construction by many-fold when testing against graphs made from assemblies produced by the HPRC (§5.5).

To support interchange with other pangenome tools or text-based processing, *odgi view* converts a graph in ODGI binary format to GFAv1. ODGI utilizes the PanSN (Garrison, 2021) specification to embed sample and haplotype information in the sequence name. This harmonizes the biosample information present in FASTA, GFA, PAF, VCF, BED, BEDPE, SAM/BAM, and GFF/GTF formats related to the graph and its embedded genome sequences. By embedding all sequences into a single hierarchical namespace related to fundamental biological groupings in the input (e.g. biosample, individual, pooled group), PanSN allows us to utilize all assemblies in the pangenome as a combined reference coordinate model.

## 5 Results

Here, we apply our methods to a series of analyses, highlighting how ODGI can assist in exploring the biological features of pangenome graphs. We follow typical analyses that we have found critical to interpreting whole genome alignments represented in the variation graph model.

To simplify our exposition, we will extract small graph regions that are easy to interpret and describe. We focus on a handful of difficult loci from the human pangenome, extracting them from a prototype human pangenome graph built with the Pangenome Graph Builder pipeline (Garrison *et al*., 2021). Pangenome graphs built from hundreds of haplotype-resolved *de novo* genome assemblies are very large, but it is often only necessary to work with only a small portion of the genomes represented, such as a specific locus (Fig. 2a) or a smaller region (Fig. 2b to 2g), or even a single gene (Fig. 3). This simplifies the downstream analyses and reduces the resources to work only with the extracted graphs. More on graph extraction and edit operations can be found at *§*S.3.

**Fig. 2:**
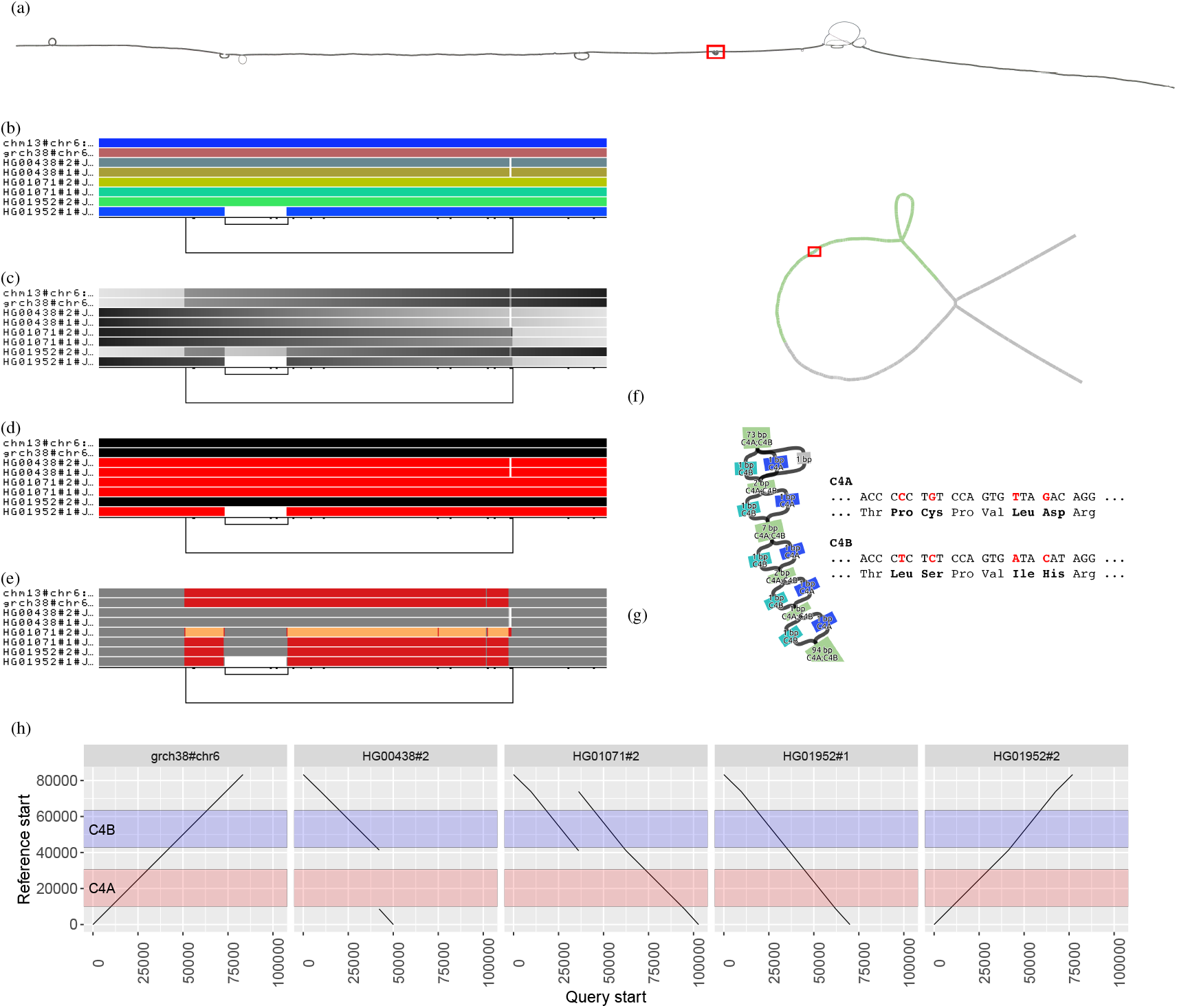
Visualizing the major histocompatibility complex (MHC) and complement component 4 (C4) pangenome graphs. **(a)** *odgi draw* layout of the MHC pangenome graph extracted from a whole human pangenome graph of 90 haplotypes. The red rectangle highlights the C4 region. **(b-e)** *odgi viz* visualizations of the C4 pangenome graph, where 8 paths are displayed: 2 reference genomes (CHM13 and GRCh38 on the top) and 6 haplotypes of 3 diploid individuals. **(b)** *odgi viz* default modality: the image shows a quite linear graph. The links at the bottom indicate the presence of a structural variant (long link) with another structural variant nested inside it (short link on the left). **(c)** Color by path position. The top two reference genomes and one haplotypes (HG01952#2) go from left to right, while 5 haplotypes go in the opposite direction, as indicated by the black color on their left. **(d)** *odgi viz* color by strandness: the red paths indicate the haplotypes that were assembled in reverse with respect to the 2 reference genomes. **(e)** *odgi viz* color by node depth: using the Spectra color palette with 4 levels of node depths, white indicates no depth, while grey, red, and yellow indicate depth 1, 2, and greater than or equal to 3, respectively. Coloring by node depth, we can see that the two references present two different allele copies of the C4 genes, both of them including the HERV sequence. The entirely grey paths have one copy of these genes. HG01071#2 presents 3 copies of the *locus* (orange), of which one contains the HERV sequence (gray in the middle of the orange). In HG01952#1, the HERV sequence is absent. **(f)** Layout of the C4 pangenome graph made with the *Bandage* tool (Wick *et al*., 2015) and annotated by using *odgi position*. Green nodes indicate the C4 genes (in red). The red rectangle highlights the regions where *C4A* and *C4B* genes differ. **(g)** Annotated *Bandage* layout of the C4 region where *C4A* and *C4B* genes differ due to single nucleotide variants leading to changes in the encoded protein sequences. Node labels were annoted by using *odgi position*. **(h)** Visualization of *odgi untangle* output in the C4 pangenome graph: the plots show the copy number status of the sequences in the C4 region with respect to the GRCh38 reference sequence, making clear, for example, that in HG00438#2, the *C4A* gene is missing (no black lines in the region annotated in red).

**Fig. 3:**
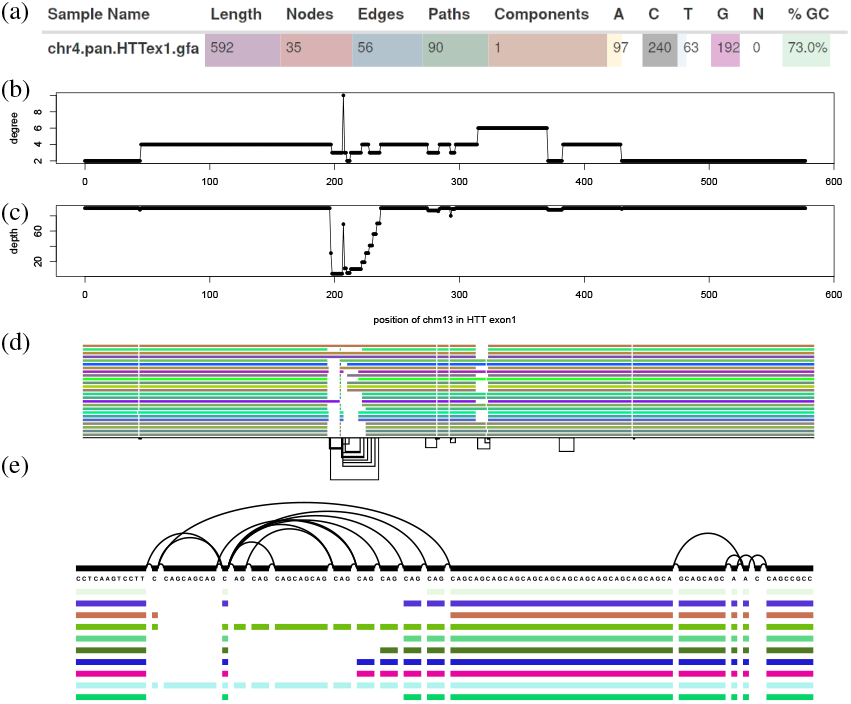
Features of a 90-haplotype human pangenome graph of the exon 1 huntingtin gene (*HTTexon1*): **(a)** Excerpt of vital statistics of the *HTTexon1* graph displayed by MultiQC’s ODGI module. **(b)** Per nucleotide node degree distribution of CHM13 in the *HTTexon1* graph. Around position 200 there is a huge variation in node degree. **(c)** Per nucleotide node depth distribution of CHM13 in the *HTTexon1* graph. The alternating depth around position 200 indicates polymorphic variation complementing the above node degree analysis. **(d)** *odgi viz* visualization of the 23 largest gene alleles, CHM13, and GRCh38 of the *HTTexon1* graph. **(e)** *vg viz* nucleotide-level visualization of 10 gene alleles, CHM13, GRCH38 of the *HTTexon1* graph focusing on the CAG variable repeat region.

### 5.1 Visualizing pangenome graphs

Visualization methods help us quickly gain insight into otherwise opaque biological data. We find visualization essential for understanding pangenome graphs. We pursue a novel approach to visualization with *odgi draw* and *odgi viz*, two tools that provide scalable ways of generating raster images showing the high-level structure of even large pangenome graphs (Fig. 2).

Using *odgi extract*, we extracted the major histocompatibility complex (MHC) locus from a 90-haplotype human chromosome 6 pangenome graph from the HPRC. Specifically, the graph contains the human references GRCh38, CHM13, and the contigs of 44 diploid individuals that encode all possible variations including those in telomeres and centromeres. The MHC genes are involved in antigen presentation, inflammation regulation, the complement system, and the innate and adaptive immune responses (Shiina *et al*., 2009). MHC genes are highly polymorphic, i.e., there are multiple different alleles across individuals in a population. Such variability becomes evident when we apply *odgi draw* to visualize the graph layout of a human MHC pangenome graph (Fig. 2a) (of note, *odgi layout* first generates the drawn projection, see §5.3). The visualization displays the graph topology in 2 dimensions (2D), with structural variation that appears as bubbles in the layout. A 2D rendering can be costly to compute, but we provide an implementation that scales linearly with pangenome sequence size, allowing us to apply it to large pangenome graphs.

The MHC locus includes the complement component 4 (C4) region, which encodes proteins involved in the complement system. In Fig. 2a, C4 corresponds to the small bubble highlighted by the red rectangle. As an example use case, we took a closer look at the C4 region of the MHC by extracting it from the full MHC pangenome graph with *odgi extract*. Then, we visualized this subgraph by applying *odgi viz*, which produces binned, linearized renderings in 1 dimension (1D), where the graph is ordered in 1D across the horizontal axis, with each path represented by a row of the vertical axis (Fig. 2b to 2e). For each path, graph nodes are arranged from left to right, with the colored bars indicating the paths and the nodes they cross. White spaces indicate where paths do not traverse the nodes. Directly consecutive nodes are displayed with no white space between the two. The meaning of the colors depends on how *odgi viz* is executed. By default, path colors are derived from the path names (Fig. 2b), which are displayed on the left of the paths. The black lines on the bottom indicate the edges connecting the nodes and, therefore, represent the graph topology (see §S.1 for a more detailed explanation). This visualization is computed in linear-time and offers a human-interpretable format suitable for understanding the topology and genome relationships in the pangenome graph. In humans, the C4 gene exists as 2 functionally distinct genes, *C4A* and *C4B*, which both vary in structure and copy number (Sekar *et al*., 2016). In combination with the observed changes in path self-coverage, which represents copy number of a given path relative to the graph (Fig. 2e), the longer link at the bottom of Fig. 2b to 2e indicates that the copy number status of these genes varies across the haplotypes represented in the pangenome. Moreover, the short nested variation on the left of the locus highlights that *C4A* and *C4B* genes segregate in both long and short genomic forms, distinguished by the presence or absence of a human endogenous retroviral (HERV) sequence.

Nevertheless, complex, nonlinear graph structures are difficult to interpret in a low number of dimensions. To overcome this limitation, *odgi viz* supports multiple visualization modalities (Fig. 2c to 2e), making it easy to grasp the properties and shape of the graph. For example, we can color the paths by path position (Fig. 2c), with light grey indicating where paths begin and dark grey where they end. This visualization is suitable for understanding graph node order, as smooth color gradients indicate that the node order respects the linear paths’ coordinate systems. Pangenome graphs can represent both strands of the genomic sequences of the DNA. We can display such information by coloring the paths by orientation, with paths colored where their sequence is reverse-complemented (red) or in direct orientation (black) with respect to the sequences of the graph nodes (Fig. 2d). Furthermore, we can use multiple color palettes to color the paths by how many times they traverse a node, which can be referred to as the path’s depth or coverage of the node, the node depth. This highlights that in the C4 pangenome graph, the haplotypes present different number of copies of the C4 genes (Fig. 2e).

### 5.2 Untangling and navigating the pangenome

The key data in a pangenome graph is a representation of the alignment (i.e., the homology relationships) between genomic sequences. Navigating and understanding the graph requires coordinate systems to link other data to the sequences represented in the graph model. ODGI’s tools use the embedded sequences to provide a universal coordinate space that is graph-independent, thereby remaining stable across different graphs built with the same sequences. Such a a universal coordinate system allows us to support several kinds of “lift-over” of coordinates between different sequences in the same or different graphs. As a demonstration, we took the C4 pangenome graph and added to its nodes gene annotation from GRCh38 (in GFF format file) using *odgi position* (§S.2.1). The resulting TSV contains pairs of nodes and colors. Taking the graph and the TSV into Bandage (Wick *et al*., 2015), the actual C4 genes are highlighted (Fig. 2f). Zooming to the nucleotide level, the annotation shows the single nucleotide differences of the *C4A* and the *C4B* genes (Fig. 2g).

*odgi position* can also translate graph and path positions between or within graphs, emitting the liftovers in BED format. For a precise translation process when conversing a query position to a reference position in a repeat region, we apply the *path jaccard* context mapping concept. It could be that the found reference node is visited several times by the reference. To ensure a precise translation, we select the reference position whose context (the multiset of *Node*.*id*s reached within a distance of e.g. 10kbp) has the best jaccard metric when compared to the query context. For a more detailed explanation of the *path jaccard* concept see §S.2.2.

To obtain a more precise overview of the locus in Fig. 2b to 2e, we applied *odgi untangle* with GRCh38 as a reference. *odgi untangle* segments paths into linear segments by breaking these segments where the paths loop back on themselves. In this way, we obtain information on the position and copy number status of the sequences in the collapsed locus, in BEDPE or PAF format. In the representation in Fig. 2h, the orientation of the line indicates if the copy number is in forward or in reverse orientation compared to GRCh38. *odgi untangle* is able to work with any sets of reference sequences, converting the graph to lift-over maps compatible with standard software for projecting annotations and alignments from one genome to another. An explanation of the untangling process is given in §S.2.

### 5.3 Latent graph structure reveals underlying biology

Pangenome graphs can hide their underlying latent structures, introducing difficulties in the analysis and interpretation. Among the causes of this is the correct ordering of the graph nodes in a convenient number of dimensions. ODGI provides a variety of sorting algorithms to find the best graph node order in 1 or 2 dimensions, allowing us to understand the sparse structures typically found in pangenome graphs and the genetic variation they represent. *odgi sort* allows the chaining of these sorting algorithms. As many of the algorithms are affected by the initial node order, this allows us to generate sorting pipelines that progressively refine the graph ordering.

We applied several of *odgi sort*’s 1D algorithms to a 90-haplotype human MHC pangenome and a C4 subgraph (Fig. S2). The randomly sorted MHC graph (Fig. S2a) hides its linear graph structure, whereas our novel path-guided (PG) stochastic gradient descent (SGD) algorithm, PG-SGD, is able to produce a globally linear ordered graph revealing the C4 region (Fig. S2b). This exploits path information to order the graph nodes. PG-SGD learns a 1D or 2D organization of the graph nodes that matches nucleotide distances in graph paths (i.e., the sequences embedded in the graph). To scale to large graphs, we learn this projection in parallel via a HOGWILD! approach (Niu *et al*., 2011). PG-SGD can be seen as an adaptation of SGD-based drawing (Zheng *et al*., 2018) to pangenome graphs. In parallel, each HOGWILD! thread updates the relative position of pairs of nodes so that their distance in the layout, or their order, better-matches their nucleotide distance in the paths running through the graph. Following standard SGD approaches, the learning rate is reduced as the algorithm progresses, and execution continues until the adjustments to the model fall below a target threshold *E*.

A PG-SGD sorting of C4 compresses both sides of the variant bubble into one dimension, leading to an interrupted pattern of nodes across the copy-number variable region (Fig. S2c). Subsequently applying a topological sort clarifies the graph’s latent structure, simplifying interpretation (Fig. S2d). To find the best order of graph nodes in 1D, *odgi sort*’s multiple sorting algorithms can be combined into a sorting pipeline to take advantage of the strength of each (results not shown). ODGI can project vector (in 1D) and matrix (2D) representations of the graph relative to these learned coordinate spaces. Based on this projection, we can trivially sort graph nodes in 1D. Moreover, we support the same concept in 2D in *odgi layout* by providing a 2D implementation of the PG-SGD algorithm (Figure 2a). A detailed description of the node ordering process can be found at §S.4. As we have shown above, the node order is crucial to understand the biological features of a pangenome graph.

### 5.4 Graph features highlight variation

Graphs statistics provide alternative ways to gain insight into pangenomes complexity revealing the overall structure, size, and features of a graph and its sequences.

As a use case study (Fig. 3), we took a look at the metrics of a 90-haplotype human pangenome graph of the exon 1 huntingtin gene (*HTTexon1*). In particular, we obtained the number of nodes, edges, paths, components, bases, the graph length, and the GC content with *odgi stats*. The output pangenome statistics in YAML textual file format was given to MultiQC’s (Ewels *et al*., 2016) newly added ODGI module. As can be seen in Figure 3a, we observe a very high GC content of 73.0% in the *HTTexon1* graph compared to the human genomic mean GC content of 40.9% (Piovesan *et al*., 2019). This is in accordance with the literature (Neueder *et al*., 2017). Despite this discovery, the MultiQC module provides an interactive way to comparatively explore statistics of an arbitrary number of graphs.

To investigate in detail which intricate regions in the *HTTexon1* graph are responsible for its genetic variation and high GC content, we took a look at the per nucleotide node degree (Fig. 3b) and node depth (Fig. 3c) distributions of CHM13 by using *odgi depth*’s and *odgi degree*’s BED output, respectively. The results indicate a highly polymorphic region around position 200 in the graph. Figure 3d supports this analysis. Zooming in on this region with *vg viz*, we can clearly identify the typical *HTTexon1* CAG variable repeat region (Fig. 3e). Figures 3b-3d highlight the variant region around position 200 of CHM13, showing the variable number of glutamine residues of the different individuals as reported by (Nance *et al*., 1999).

### 5.5 Performance evaluation

Although many of the operations that ODGI provides are unique, some are common with the existing VG toolkit. We compare with these to highlight the practical performance implications of our graph data structure design. Our results highlight the efficient parallel algorithm implementations enabled by this design.

We compared the efficiency of ODGI (v0.6.3-56-gebc493f “Pulizia”) and VG (v1.37.0 “Monchio”) for routine pangenome tasks. In particular, we measured the execution time and memory usage (i) of transforming a GFAv1 file into a tool’s native format, (ii) the extraction of a subgraph, (iii) the visualization of a pangenome graph, and (iv) the finding of path positions in a pangenome graph. These graph operations are key when it comes to the understanding of pangenome graphs. They are also a set of functions implemented in both toolkits. We ran these operations for a varying number of threads and haplotypes in the graph for a scaling analysis. We ran each evaluation configuration 10 times and report the mean of each run. All evaluations were performed on a VM in the German Network for Bioinformatics Infrastructure (deNBI) cloud with 28 cores and 256GB of RAM. The presented results are from a 90-haplotype chromosome 6 human pangenome graph built with data from the HPRC. Specifically, the graph contains the human references GRCh38, CHM13, and the contigs of 44 diploid individuals that encode all possible variations including those in telomeres and centromeres. When transforming a GFAv1 file with VG, the static XG file format was used. The tools involved in the evaluation process require the XG format.

In general, ODGI makes comparatively better use of multithreading and requires much less memory (Fig. 4, Tab. S4) across all operations. ODGI scales much butter than VG when working with complex regions of the graph. For example, extracting a difficult centromeric subgraph (Fig. 4b), ODGI is up to 40 times faster and requires 8 times less memory than VG.

**Fig. 4:**
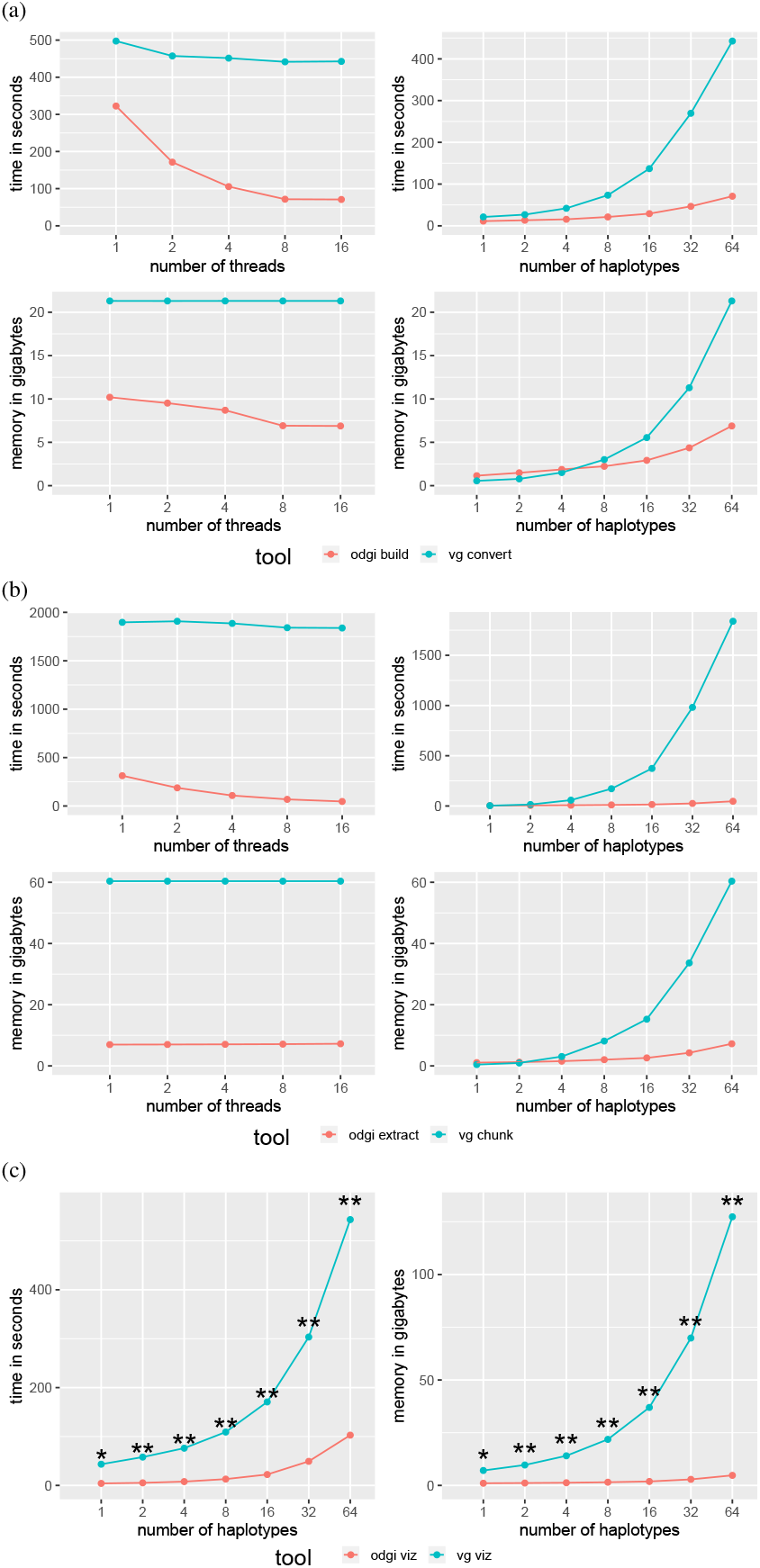
Performance on a graph of human chromosome 6 from the HPRC. ODGI compares favorably to VG across all routine pangenomic tasks. Evaluations across threads were done using a 64 human haplotype graph. Evaluations across haplotypes were done using 16 threads. **(a)** Performance evaluation when translating a graph into the tools’ respective native formats. **(b)** Performance evaluation when extracting the centromeric region from the HPRC graph. **(c)** Performance evaluation when visualizing a graph. Both tools were run with only one thread. *vg viz*: **\***A 816MB SVG was produced which can’t be opened by any program. **\*\***All produced SVGs only contain an XML header, nothing else.

Both visualization tools can only make use of a single thread. For a 1 haplotype, graph *vg viz* produces a 816MB SVG which can’t be opened by the standard programs to date. For larger graphs, *vg viz* runs through and produces SVGs with only the XML header. This makes it unusable for large graphs.

We also measured the disk space usage of GFAv1, ODGI’s, and VG’s binary formats (Tab. S5). While VG’s XG occupies less disk space for smaller graphs, ODGI requires less space for graphs having 32 haplotypes or more. We hypothesize that this indicates the lower marginal cost for additional haplotypes when using ODGI’s id delta encoding scheme.

## 6 Discussion

Pangenome graphs stand to become a ubiquitous model in genomics thanks to their capability to represent any genetic variant without being affected by reference bias (Eizenga *et al*., 2020b). However, despite this great potential, their spread is impeded by the lack of tools capable of managing and analyzing pangenome graphs easily and efficiently.

By providing a set of standard analysis “verbs” to interact with pangenome graphs, ODGI enables users to explore and discover important biological features captured in this flexible, inclusive model. It provides tools to easily transform, analyze, simplify, validate, and visualize pangenome graphs at large scale. In particular, lifting over annotations and linearizing nested graph structures place the suite as the bridge between traditional linear reference genome analysis and pangenome graphs. With the increased adoption of long read sequencing we expect pangenomic tools to become increasingly common in the genomic studies at different taxonomic levels and in biomedical research. This progression is already afoot, particularly for targets that involve complex variation, such as cancer (The Computational Pan-Genomics Consortium, 2016), plant pangenomics (Bayer *et al*., 2020; Liu *et al*., 2020; Qin *et al*., 2021; Li *et al*., 2022; Bayer *et al*., 2022), and metagenomics (Zhong *et al*., 2021). Also, when studying animals like bovines (Leonard *et al*., 2021; Talenti *et al*., 2022; Bovine Pan-Genome Consortium, 2022).

Currently, bacterial pangenomes are best handled by specialized tools like PPanGGolin (Gautreau *et al*., 2020), PanGraph (Noll *et al*., 2022) or PanX (Ding *et al*., 2017). The latter one doesn’t build a graphical representation of a pangenome. But, it already has a very developed ecosystem, which allows a detailed analysis of bacterial pangenomes using an interactive GUI. Unlike these approaches, which provide a monolithic, integrated solution to understanding pangenomes, ODGI is designed as a low-level toolkit that can work on a generic pangenome graph model frequently used by other existing methods. We hope that this design renders it useful to pangenome analysis pipeline authors. Other pangenome analysis platforms, like PanTools (Sheikhizadeh *et al*., 2016) provide access to pangenome analyses at the scales we demonstrate with ODGI, but use specialized de Bruijn graph models to achieve this. In contrast ODGI supports the highly generic variation graph model, which has greater representational power than de Bruijn graphs.

ODGI will facilitate disentangling, describing and analyzing a much larger set of variation than previously was possible with tools that depend on short reads and reference genomes. Furthermore, users can even consider ODGI as a framework, taking advantage of its algorithms to develop new and more advanced tools that work on pangenome graphs, thus expanding the type of possible pangenomic analyses available to the scientific community.

The performance analysis shows that ODGI outperforms VG when handling large, complex pangenome graphs. Across the evaluation of key graph operations, ODGI’s memory peak was 10GB. This makes it perfectly suited to be run interactively on a recent laptop. We expect that ODGI will be able to handle the next phase of the HPRC, a pangenome graph constructed from 300 individuals, without any problems.

While ODGI does not construct graphs from scratch nor is is capable of extending them, it is already the backbone of the Pangenome Graph Builder pipeline (Garrison *et al*., 2021). Its static, large-scale 1D and 2D visualizations of the pangenome graphs allow an unprecedented high-level perspective on variation in pangenomes, and have also been critical in the development of pangenome graph building methods. However, an interactive solution that combines the 1D and 2D layout of a graph with annotation and read mapping information across different zoom levels is still missing. Recent interactive pangenome graph browsers are reference-centric (Beyer *et al*., 2019; Yokoyama *et al*., 2019), have a limited predefined coordinate system (Durant *et al*., 2021), or focus primarily on 2D representations (Wick *et al*., 2015; Gonnella *et al*., 2018). Our graph sorting and layout algorithms can provide the foundation for future tools of this type. We plan to focus on using these learned models to detect structural variation and assembly errors.

ODGI has allowed us to explore *context mapping* deconvolution of pangenome graph structures via the path jaccard metric. This resolves a major conceptual issue that has strongly guided existing algorithms to construct pangenome graphs. Previously, great efforts have been made to prevent the “collapse” of non-orthologous sequences in the graph topology itself (Li *et al*., 2020). This has been seen as essential to making these new bioinformatic models interpretable. While our presentation is primarily qualitative, our work demonstrates that we can mitigate this issue by exploiting the pangenome graph not as a static reference, but as a dynamic model of the mutual alignment of many genomic sequences. Because pangenome graphs can contain complete genomes, we are able to query them to polarize the information they contain in easily-interpretable and reusable pairwise formats that are widely supported in bioinformatics. ODGI also projects variation graphs into vector and matrix representations that allow the direct application of machine learning and statistical models to the pangenome. We expect that ODGI will provide a reference interface between pangenomic and genomic approaches for understanding genome variation.

## Acknowledgments

We thank members of the HPRC Pangenome Working Group for their insightful discussion and feedback, and members of the HPRC production teams for their development of resources used in our exposition.

## Data availability

Code and links to data resources used to build this manuscript and its figures, can be found in the paper’s public repository: https://github.com/pangenome/odgi-paper.

## S Supplement

### S.1 Graph topology visualization

In an *odgi viz* visualization, given a path X traversing two nodes A and B, the corresponding edge is represented by a black line starting from the left or right of node A if this node is traversed in reverse or forward, respectively, by path X; the black line ends to the left or right of node B if this node is traversed in forward or reverse, respectively, by path X. Consequently, if two consecutive nodes are linked both in forward, no edge is shown (it would be 0 pixels long, as it would start from the right of the first node and ends to the left of the second one).

### S.2 Graph navigation and untangling

Pangenome graphs can model alignments of many genomes. With *odgi untangle*, users can extract pairwise alignment information between a given set of “query” sequences and a given set of “target” sequences (used as references). While pangenome graphs may contain looping structures that imply many-to-many alignments between the pangenome sequences, these untangled alignments map each segment of the queries to a single segment in the set of targets. *odgi untangle* first discovers segment boundaries using standard approaches for detecting repeats in sequence graphs (Pevzner, 2004). We finally “untangle” by finding the target segment that best matches each query segment using the *path jaccard* context mapping model. Moreover, to obtain base-level precise information on the relationships between the repeated sequences, we can align them by using the pairs of regions that came from the untangling to guide the alignment (Guarracino *et al*., 2021). *odgi tips* can identify the break point positions of the contigs relative to the reference(s) in the graph by walking from the ends of each contig until a reference node is found. It could be that the reference visits the node several times. Therefore, for each contig range (a tip) *odgi tips* takes a look at each possible reference window and finds the most similar one using the *path jaccard* concept. The output is a BED file with the best reference hit and position for each of the contigs’ ends.

#### S.2.1 The GFF liftover

A GFF file contains annotations for one or more paths in the graph. For each annotation, we know the start and end within that path. So we can annotate all nodes that are visited by such a path range with the information from the attribute field. If there are overlapping features, we append the annotation for each node. Using the same coloring schema as in *odgi viz* we generate a color for each annotated node by its collected annotation.

If a subgraph was as a result from e.g. *odgi extract*, the path names are usually in the form of name:start-end. *odgi position* is able to automatically detect this and adjust the positions given in the GFF on the fly to the new positions given in the subgraph. For each GFF entry, it just subtracts the “missing” number of nucleotides from the start and end field. That’s how we adjust for the subgraph annotation.

#### S.2.2 Path Jaccard concept

*odgi tips, odgi untangle*, and *odgi position* all translate a query path position to a target path position. In a repeat region, it could be that the node of the query path position (*N*_*q*_) is visited several times by the target path. We use the graph *path jaccard* concept to infer the best target position for a given query path position. For the query position we look at each possible target window and find the most-similar one:

1. Starting from *N*_*q*_ we follow the steps of one target path position a certain nucleotide distance to the left and a certain nucleotide distance to the right, also called context. The ODGI default distance is 10kbp for each direction.
2. We collect all node identifiers that are visited this way in a multi-set.
3. Repeat the same process when following the query path position steps.

Now we have two multi-sets of node identifiers, the target path position *T* one, and the query path position *Q* one. We apply the Jaccard measure to estimate the similarity between these two multi-sets (Equ. 1).

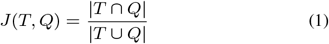

The “|” indicate that, after we joined our sets together, we actually calculate the total length in nucleotides represented by the intersection node set in the nominator, and by the union node set in the denominator, respectively. The division of the two nucleotide lengths then gives us the jaccard measure for the two path positions. We collect these jaccard measurements for all possible target path positions. The largest *path jaccard* determines the actual target path position for the translation.

Dependent on how repeat-streaked the graph is and how it was constructed, one might want to adjust context size, to get an even more precise positional translation. If we can’t follow the full distance to one direction, because we are at an end of a sequence, we also adjust this for all other context evaluating windows.

In the following an example which is based on the graph in Figure S1. Let’s assume, we want to translate the query path position *query:3* to a path position in path *target* with a context distance of 2 nucleotides per direction. Node *2* is where *query:3* is located. The node is traversed two times by *target*. Therefore, we apply the *path jaccard* concept to do a precise position translation. Starting from node *2*, we collect nodes {1, 2, 4} for path *query*. For path *target* we have two possible steps to start our collection from: The first or the second step in node *2*. We obtain the node sets *T*_1_ = {1, 2, 2} and *T*_2_ = {2, 2, 3}.

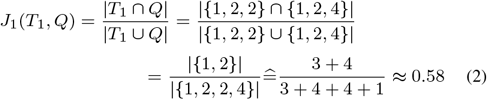

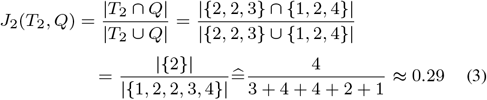

We observe a *path jaccard* of 0.58 for *T*_1_*/Q* (Equ. 2) and 0.29 for *T*_2_*/Q* (Equ. 3). Therefore, we translate query path position *query:3* to the target path position *target:3*.

### S.3 Editing

Subgraphs can be extracted by using the paths in the graph as coordinate systems to guide the process. For such operation, *odgi extract* allows users to extract specific regions of the graph as defined by query criteria. Regions of interest can be specified by graph nodes or path range(s), also in BED format. Furthermore, it is possible to indicate a list of paths to be preserved completely in the extracted graph. We begin by collecting all graph nodes that fall within the ranges to extract (and the paths to preserve, if requested). Users can specify the number of steps or nucleotides to expand the selection and include neighboring nodes. Then, edges connecting all selected nodes are added in the subgraph under construction. Finally, the portions of the paths (i.e., the subpaths) walking through the selected nodes are extracted and added to the new subgraph. Subpaths are searched in parallel, created serially, and extended in parallel again thanks to the parallelism enabled by the ODGI data structure (see §4.1), making *odgi extract* a scalable solution to extract also complex subregions presenting nodes with very high node depth.

Pangenome graphs can embed multiple chromosomes as separated connected components (inter-chromosomal structural variants would join the components into bigger ones). *odgi explode* separates the connected components in different ODGI format files, while *odgi squeeze* allows merging multiple graphs into the same ODGI format file, preventing node ID collisions.

Pangenome graphs can be used in a variety of applications, ranging from read mapping to variant identification and genotyping (Eizenga *et al*., 2020b). However, graphs presenting complex topology can increase the computational overhead of many downstream analyses. ODGI offers multiple commonly-needed basic operations on the topology of the graph and its nodes.

For simplifying the graph structure, users can use *odgi prune* to take away complex parts as defined by query criteria, while with *odgi break* they can remove cycles in the graph, reducing the complexity of the graph topology. Furthermore, *odgi groom* allows removing spurious inverting links by exploring the graph from the orientation supported by most paths; the process does not remove any genetic information, but only edits how the sequences are represented in the graph.

To enable efficient sequence alignment against the graph, long nodes can be divided into shorter nodes at a maximum requested size using *odgi chop*. Partial order alignment, which consists of aligning sequences against a directed acyclic graph (DAG), is frequently used in pangenome building pipelines (Garrison *et al*., 2021), but the current implementations return DAGs with 1-bp long nodes. *odgi unchop* allows joining nodes that can be merged without changing the graph topology, nor the embedded sequences, obtaining an equivalent, but more compact, representation of the graph.

### S.4 Sorting and node identifier compaction

Most subcommands in ODGI require and verify that the input graph’s node identifiers (IDs) are optimized, that is compacted from 1 to N where N is the number of nodes in the graph. If this assumption is violated, *odgi sort* provides functionality to optimize the graph. This means that the first node identifier (ID) starts at 1 and the last node ID is the number of nodes. All sorting operations update the graph in place with an efficient ID rewriting algorithm. The graph is then updated in place. First, the node identifiers are normalized (from 1 to number of nodes) including the adjustment of the edges. Second, path information, including both path metadata that points into the start and end steps of the path, plus each step of every path, is updated, too. We point out that changing the node order does not change our coordinate systems based on paths. These will now refer to a new node ordering.

When we sort a graph, we switch the node IDs of the nodes according to the result of the sorting algorithm. For example, if a random sort was applied, all existing node IDs would be replaced with new, random ones (the largest node ID would still correspond to the number of nodes on the graph). We would update the edge and path information as described in the paragraph above. The reordering of nodes has a great influence on how the pangenome looks like (Fig. S2).

### S.5 Linear projections

Pangenome graph topology can be derived by applying *odgi matrix*, obtaining information on graph nodes connections in textual sparse matrix format. To investigate on the genomic sequences encoded in the graph, *odgi paths* allows users to calculate pairwise overlap statistics of groupings of paths and emit all path sequences in FASTA format, and it also allows the generation of a “pangenome matrix” that reports the copy number (presence/absence) of each path over each node.

In standard practice, pangenome analysis examines presence-absence variations (PAVs), which correspond to loci that are present in some samples but not others. *odgi pav* uses a set of genomic intervals (in BED expressed in the coordinate space of paths in the graph) to demarcate PAVs. It then describes the coverage of all paths relative to these PAVs, yielding matrix or tabular representation. The precise determination of PAVs based on graph topology remains an open problem, but practically any method capable of generating a BED file can be used here. This lets us define PAVs using repeat or gene annotations of the genomes represented as paths in the graph. A simple technique is to take the output of *odgi flatten*, which generates a linearization of the graph by emitting the pangenome sequence (the concatenation of all node sequences) in FASTA format, and the projection of all paths on the linearized sequence in BED format relative to the graph’s paths.

ODGI also supports an experimental binned representation of graphs designed to support the study and visualization of pangenomes at different scales of resolution. *odgi bin* summarizes graph path information into bins of a specified size, generating a summarized view of gigabase scale graphs in TSV or JSON file formats. We have further supported this binning approach in pangenome graph ontology model (Yokoyama *et al*., 2020).

### S.6 Evaluation

**Fig. S1:**
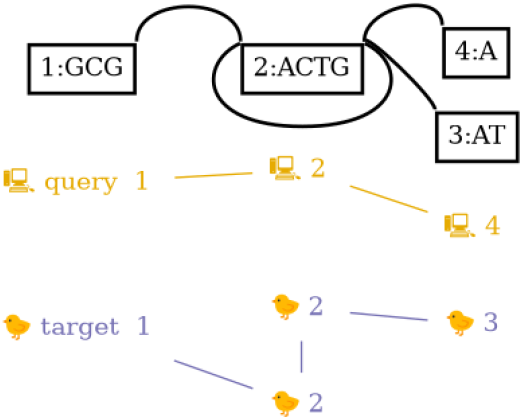
Simple example graph to demonstrate the *path jaccard* concept. Generated with *vg view* and *graphviz’ dot* command. On top are the nodes connected with edges. On the bottom the paths through the nodes. Each path has a different color and emoji.

**Fig. S2:**
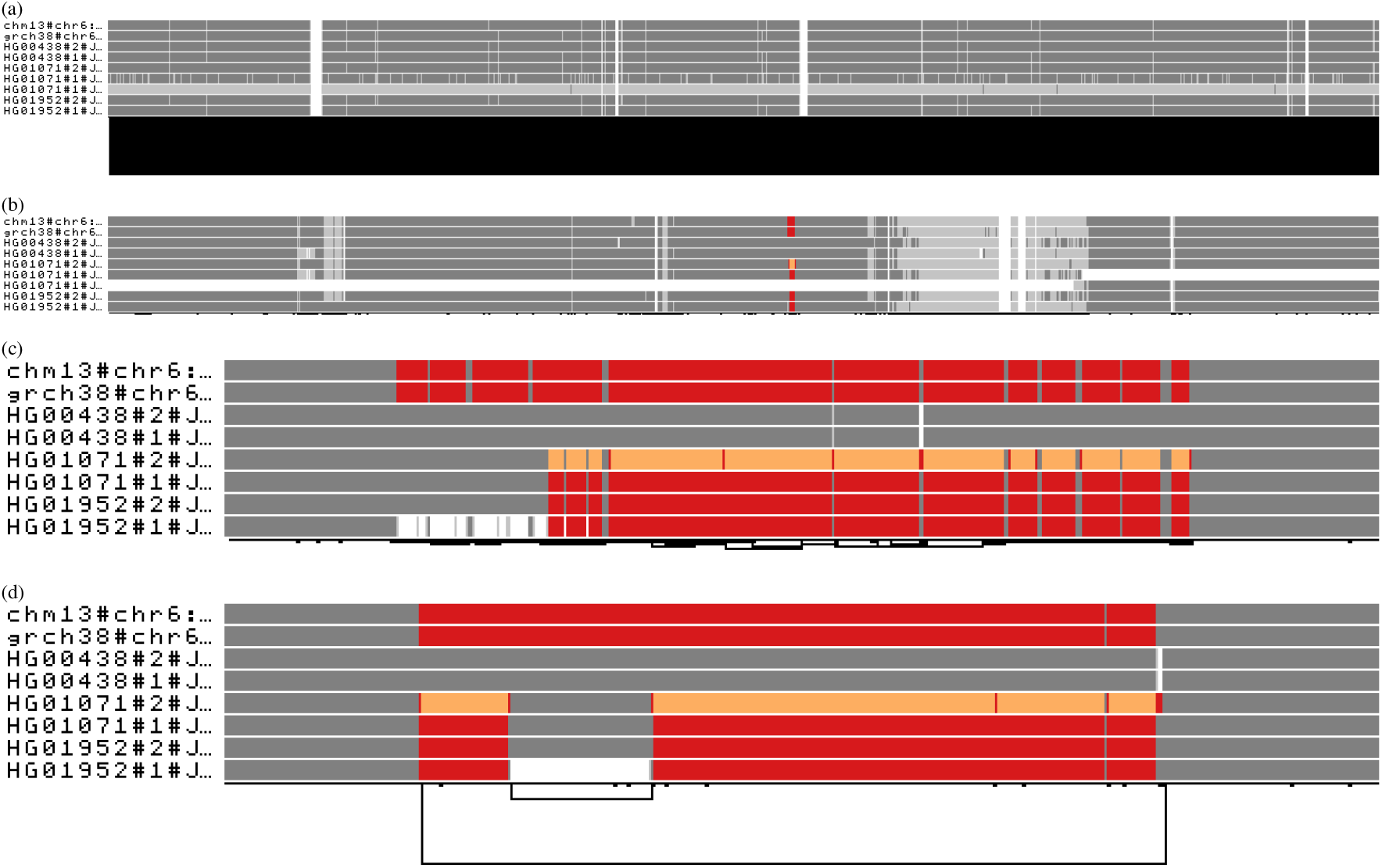
Sorting and visualizing the human major histocompatibility complex (MHC) and complement component 4 (C4) pangenome graphs. All visualization are made with *odgi viz* by coloring the paths by node depth: using the Spectra color palette with 4 levels of node depths, white indicates no depth, while grey, red, and yellow indicate depth 1, 2, and greater than or equal to 3, respectively. **(a)** Visualization of a MHC pangenome graph with nodes randomly sorted from left to right. The image shows a region 5 Mbp-long. The bad node order completely hides the linear graph structure. **(b)** Visualization of the same MHC pangenome graph but sorted by applying the path-guided stochastic gradient descent algorithm (PG-SGD). The graph globally shows a linear structure, without long range links (at the bottom of the image). Furthermore, the C4 region is visible as a region with red and orange paths. **(c)** Visualization of the PG-SGD sorted C4 subgraph. The image shows a region 83361 bp-long. The C4 region is still not well sorted locally, with the node order that still hides the underlying copy number variation status present in such a region. **(d)** Visualization of the same C4 pangenome graph but sorted by applying a topological sorting. The graph shows its structure: the two references present two different allele copies of the C4 genes (in red), both of them including the HERV sequence. HG01071#2 presents 3 copies of the *locus* (orange), of which one contains the HERV sequence (gray in the middle of the orange). In HG01952#1, the HERV sequence is absent (white in the middle of the red).

**Table S1:** Performance measurements when transforming a human chromosome 6 pangenome graph into the tool’s native format. **haps** is the number of haplotypes in the graph. Displayed are the mean results after 10 runs.

| threads | haps | time in seconds |  | memory in gigabytes |  |
| --- | --- | --- | --- | --- | --- |
|  |  | odgi build | vg convert | odgi build | vg convert |
| 1 | 1 | **10.78** | 21.92 | 1.15 | **0.56** |
| 1 | 2 | **14.55** | 28.71 | 1.48 | **0.78** |
| 1 | 4 | **22.28** | 45.02 | 1.82 | **1.51** |
| 1 | 8 | **43.41** | 78.67 | **2.52** | 3.01 |
| 1 | 16 | **76.60** | 133.27 | **3.57** | 5.54 |
| 1 | 32 | **176.55** | 292.32 | **5.90** | 11.29 |
| 1 | 64 | **322.50** | 497.51 | **10.19** | 21.3 |
| 2 | 1 | **10.56** | 21.18 | 1.15 | **0.56** |
| 2 | 2 | **12.79** | 27.51 | 1.48 | **0.78** |
| 2 | 4 | **17.08** | 43.64 | 1.81 | **1.51** |
| 2 | 8 | **28.90** | 75.52 | **2.42** | 3.01 |
| 2 | 16 | **47.93** | 126.71 | **3.46** | 5.54 |
| 2 | 32 | **102.67** | 280.12 | **5.58** | 11.29 |
| 2 | 64 | **171.31** | 457.37 | **9.52** | 21.3 |
| 4 | 1 | **10.54** | 20.69 | 1.15 | **0.56** |
| 4 | 2 | **12.63** | 27.90 | 1.48 | **0.78** |
| 4 | 4 | **14.80** | 42.36 | 1.88 | **1.51** |
| 4 | 8 | **21.00** | 74.80 | **2.18** | 3.01 |
| 4 | 16 | **32.69** | 123.41 | **3.08** | 5.54 |
| 4 | 32 | **65.30** | 271.10 | **4.87** | 11.29 |
| 4 | 64 | **105.51** | 451.54 | **8.70** | 21.3 |
| 8 | 1 | **11.53** | 20.33 | 1.15 | **0.56** |
| 8 | 2 | **13.16** | 27.18 | 1.48 | **0.78** |
| 8 | 4 | **15.61** | 41.77 | 1.85 | **1.51** |
| 8 | 8 | **19.63** | 73.09 | **2.20** | 3.01 |
| 8 | 16 | **28.43** | 131.82 | **2.87** | 5.54 |
| 8 | 32 | **46.23** | 256.74 | **4.30** | 11.29 |
| 8 | 64 | **71.55** | 441.59 | **6.91** | 21.30 |
| 16 | 1 | **11.18** | 21.14 | 1.15 | **0.56** |
| 16 | 2 | **13.27** | 26.74 | 1.48 | **0.78** |
| 16 | 4 | **15.55** | 41.90 | 1.87 | **1.51** |
| 16 | 8 | **21.16** | 73.36 | **2.24** | 3.01 |
| 16 | 16 | **29.09** | 136.96 | **2.92** | 5.54 |
| 16 | 32 | **46.58** | 269.61 | **4.36** | 11.29 |
| 16 | 64 | **70.84** | 442.71 | **6.89** | 21.30 |

**Table S2:** Performance measurements when extracting the centromeric region of a human chromosome 6 pangenome graph. **haps** is the number of haplotypes in the graph. Displayed are the mean results after 10 runs.

| threads | haps | time in seconds |  | memory in gigabytes |  |
| --- | --- | --- | --- | --- | --- |
|  |  | odgi extract | vg chunk | odgi extract | vg chunk |
| 1 | 1 | 3.65 | **3.34** | 1.07 | **0.42** |
| 1 | 2 | **6.08** | 14.25 | 1.22 | **0.95** |
| 1 | 4 | **15.96** | 58.67 | **1.51** | 3.06 |
| 1 | 8 | **37.16** | 170.63 | **1.91** | 8.12 |
| 1 | 16 | **67.47** | 364.98 | **2.49** | 15.25 |
| 1 | 32 | **161.32** | 968.71 | **4.00** | 33.65 |
| 1 | 64 | **313.05** | 1897.00 | **6.96** | 60.37 |
| 2 | 1 | 3.67 | **3.33** | 1.07 | **0.42** |
| 2 | 2 | **5.50** | 13.57 | 1.22 | **0.96** |
| 2 | 4 | **12.33** | 56.44 | **1.51** | 3.06 |
| 2 | 8 | **24.06** | 170.62 | **1.92** | 8.12 |
| 2 | 16 | **43.49** | 376.67 | **2.51** | 15.25 |
| 2 | 32 | **97.03** | 987.85 | **4.03** | 33.65 |
| 2 | 64 | **187.51** | 1907.98 | **7.00** | 60.37 |
| 4 | 1 | 3.64 | **3.32** | 1.07 | **0.42** |
| 4 | 2 | **5.59** | 13.40 | 1.22 | **0.96** |
| 4 | 4 | **9.93** | 57.67 | **1.52** | 3.06 |
| 4 | 8 | **16.58** | 174.94 | **1.94** | 8.11 |
| 4 | 16 | **27.99** | 374.99 | **2.52** | 15.26 |
| 4 | 32 | **56.94** | 992.85 | **4.06** | 33.65 |
| 4 | 64 | **108.34** | 1885.91 | **7.04** | 60.37 |
| 8 | 1 | 3.61 | **3.30** | 1.07 | **0.42** |
| 8 | 2 | **5.47** | 14.01 | 1.22 | **0.96** |
| 8 | 4 | **8.43** | 58.69 | **1.53** | 3.06 |
| 8 | 8 | **12.40** | 180.51 | **1.98** | 8.11 |
| 8 | 16 | **19.11** | 379.25 | **2.55** | 15.26 |
| 8 | 32 | **36.35** | 991.09 | **4.11** | 33.65 |
| 8 | 64 | **68.54** | 1841.57 | **7.11** | 60.37 |
| 16 | 1 | 3.64 | **3.34** | 1.07 | **0.42** |
| 16 | 2 | **5.53** | 14.67 | 1.22 | **0.95** |
| 16 | 4 | **7.62** | 58.22 | **1.54** | 3.06 |
| 16 | 8 | **10.55** | 171.90 | **2.03** | 8.12 |
| 16 | 16 | **14.80** | 373.70 | **2.60** | 15.26 |
| 16 | 32 | **25.51** | 981.79 | **4.27** | 33.65 |
| 16 | 64 | **46.87** | 1838.51 | **7.23** | 60.37 |

**Table S3:** Performance measurements when visualizing a human chromosome 6 pangenome graph. **haps** is the number of haplotypes in the graph. Both *odgi viz* and *vg viz* were run with 1 thread. Displayed are the mean results after 10 runs. **\***A 816MB SVG was produced which can’t be opened by any program. **\*\***All produced SVGs were empty except for an XML header.

| haps | time in seconds |  | memory in gigabytes |  |
| --- | --- | --- | --- | --- |
|  | odgi viz | vg viz | odgi viz | vg viz |
| 1 | **4.14** | 43.40 * | **1.04** | 7.09 * |
| 2 | **5.30** | 57.94 ** | **1.14** | 9.69 ** |
| 4 | **7.85** | 76.15 ** | **1.27** | 14.05 ** |
| 8 | **12.95** | 109.16 ** | **1.53** | 21.84 ** |
| 16 | **22.33** | 170.72 ** | **1.88** | 36.97 ** |
| 32 | **49.24** | 303.40 ** | **2.87** | 69.88 ** |
| 64 | **102.82** | 543.50 ** | **4.78** | 127.36 ** |

**Table S4:** Performance measurements when locating all path positions of a node in a human chromosome 6 pangenome graph. **haps** is the number of haplotypes in the graph. Displayed are the mean results after 10 runs. The number of threads does not affect the running time or memory consumption. *vg find* had to be run for each path position. The total run time of finding all path positions of a node is reported here.

| threads | haps | time in seconds |  | memory in gigabytes |  |
| --- | --- | --- | --- | --- | --- |
|  |  | odgi position | vg find | odgi position | vg find |
| 1 | 1 | 2.58 | **0.16** | 1.01 | **0.21** |
| 1 | 2 | 3.19 | **0.39** | 1.11 | **0.26** |
| 1 | 4 | 3.52 | **1.23** | 1.23 | **0.42** |
| 1 | 8 | **3.97** | 4.28 | 1.49 | **0.74** |
| 1 | 16 | **5.07** | 14.32 | 1.81 | **1.25** |
| 1 | 32 | **7.79** | 55.61 | 2.73 | **2.46** |
| 1 | 64 | **15.60** | 198.54 | **4.51** | 4.56 |
| 2 | 1 | 2.54 | **0.16** | 1.01 | **0.21** |
| 2 | 2 | 3.17 | **0.38** | 1.11 | **0.26** |
| 2 | 4 | 3.55 | **1.24** | 1.23 | **0.42** |
| 2 | 8 | **3.95** | 4.31 | 1.49 | **0.74** |
| 2 | 16 | **5.08** | 14.10 | 1.81 | **1.25** |
| 2 | 32 | **7.76** | 55.56 | 2.73 | **2.46** |
| 2 | 64 | **15.34** | 198.56 | **4.51** | 4.56 |
| 4 | 1 | 2.67 | **0.16** | 1.01 | **0.21** |
| 4 | 2 | 3.15 | **0.38** | 1.11 | **0.26** |
| 4 | 4 | 3.55 | **1.23** | 1.23 | **0.42** |
| 4 | 8 | **3.98** | 4.31 | 1.49 | **0.74** |
| 4 | 16 | **5.06** | 14.23 | 1.81 | **1.25** |
| 4 | 32 | **7.79** | 55.66 | 2.73 | **2.46** |
| 4 | 64 | **15.41** | 196.6 | **4.51** | 4.56 |
| 8 | 1 | 2.59 | **0.16** | 1.01 | **0.21** |
| 8 | 2 | 3.20 | **0.38** | 1.11 | **0.26** |
| 8 | 4 | 3.55 | **1.25** | 1.23 | **0.42** |
| 8 | 8 | **3.99** | 4.31 | 1.49 | **0.74** |
| 8 | 16 | **5.07** | 14.20 | 1.81 | **1.25** |
| 8 | 32 | **7.70** | 55.12 | 2.73 | **2.46** |
| 8 | 64 | **15.9** | 203.02 | **4.51** | 4.56 |
| 16 | 1 | 2.62 | **0.16** | 1.01 | **0.21** |
| 16 | 2 | 3.16 | **0.38** | 1.11 | **0.26** |
| 16 | 4 | 3.55 | **1.23** | 1.23 | **0.42** |
| 16 | 8 | **3.95** | 4.26 | 1.49 | **0.74** |
| 16 | 16 | **5.1** | 14.3 | 1.81 | **1.25** |
| 16 | 32 | **7.66** | 54.78 | 2.73 | **2.46** |
| 16 | 64 | **15.5** | 202.57 | **4.51** | 4.56 |

**Table S5:** Disk space measurements of GFAv1 and the respective tools’ binary formats. **haps** is the number of haplotypes in the graph. For VG, the file size of the XG format was measured.

| disk space in megabytes |  |  |  |
| --- | --- | --- | --- |
| haps | GFAv1 | ODGI | VG |
| 1 | 283 | 604 | **189** |
| 2 | 333 | 660 | **237** |
| 4 | 432 | 745 | **394** |
| 8 | 622 | 968 | **706** |
| 16 | 947 | 1277 | **1200** |
| 32 | 1657 | **2131** | 2388 |
| 64 | 2913 | **3745** | 4426 |

